# Accurate reconstruction of dynamic gene expression and growth rate profiles from noisy measurements

**DOI:** 10.1101/2021.03.16.435606

**Authors:** Gonzalo Vidal, Carlos Vidal-Céspedes, Timothy J. Rudge

## Abstract

Cells face changing environments to which they sense and respond in complex ways, changing their rates of gene expression and growth. Measuring these dynamics is therefore essential to understanding natural and synthetic regulatory networks that give rise to functional phenotypes. However, reconstruction of gene expression and growth rate profiles from typically noisy measurements of cell populations is difficult due to the effects of noise at low cell densities among other factors. We present here a method for estimation of dynamic gene expression rates and biomass growth rates from noisy measurement data, and show that it is several times more accurate than current approaches. We applied our method to multiple promoter-reporter fusion genes. Gene expression rates of such promoter-reporter fusions are typically used as a proxy for transcription rates. However, using our method we show that fusion gene expression rate dynamics are determined at least by the promoter of interest and the downstream reporter.

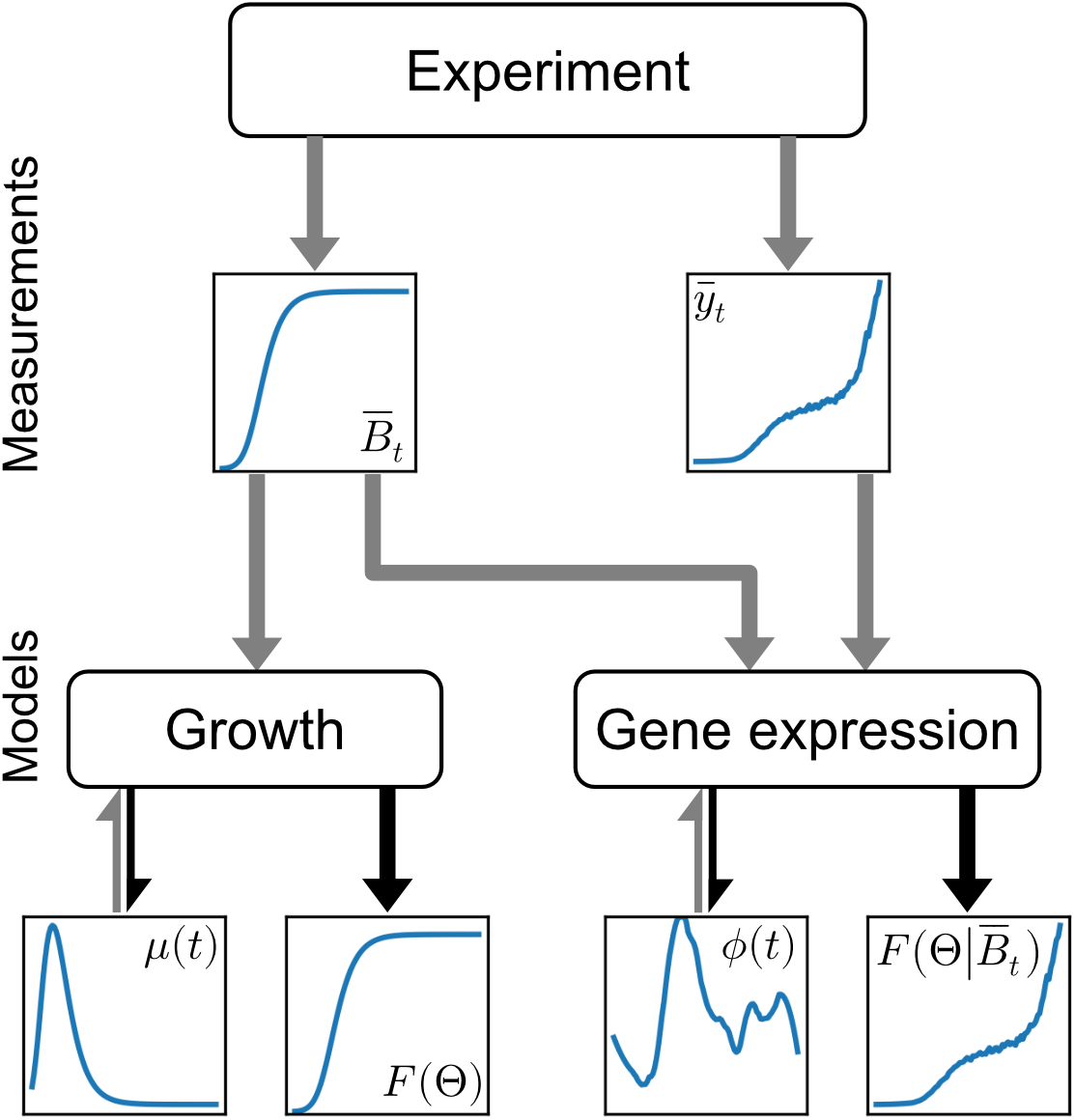

## Introduction

Measuring and analyzing the dynamics of gene expression are fundamental to understanding cellular regulation by both natural and synthetic gene networks in the face of changing growth environments. Typical experiments to measure gene expression rates in bacteria and other microorganisms utilize fluorescent reporters to track the expression levels of lineages of cells [Beal et al., 2018, Lichten et al., 2014, Kalir et al., 2001, Berthoumieux et al., 2013]. The total biomass of these lineages is also tracked, typically using optical density measurements or colony size [Beal et al., 2020, Nuñez et al., 2017]. The genes measured are often fusions of promoters of interest with a downstream fluorescent reporter, and their expression rate profiles taken to be indicative of the transcription rate of the promoter [Rudge et al., 2016, Berthoumieux et al., 2013].

Reconstructing gene expression and growth rate profiles from these data is difficult because, particularly at low biomass, measurements suffer from significant noise. Typical methods involve data smoothing and differentiation of the resulting signal [Ronen et al., 2002, Aïchaoui et al., 2012, De Jong et al., 2010]. This approach is sensitive to noise leading to the development of a more robust method based on linear inversion of differential equation models [Zulkower et al., 2015] - herein referred to as the direct method. However, as we show here, the results of this method are inaccurate particularly at low biomass. We present a method for reconstruction of gene expression and growth rate profiles that achieves several times better accuracy. Using this method on a synthetic multireporter plasmid we show that gene expression rate profiles depend on both the promoter and downstream reporter of the gene of interest.

## Results

### Gene expression and growth dynamics

A simple two-step transcription-translation model of gene expression dynamics can be formulated as [Rudge et al., 2016]:

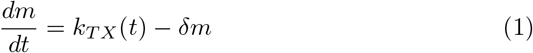

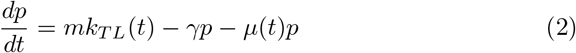

with *m* the mRNA concentration, *p* the reporter protein concentration, *k_TX_*(*t*) and *k_TL_*(*t*) the transcription and translation rates, *δ* and *γ* the corresponding degradation rates of mRNA and protein, and *μ*(*t*) is the instantaneous relative growth rate. In the typical case of short half-life mRNAs, we may assume quasi-steady state and,

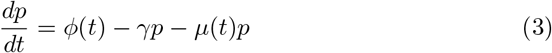

with,

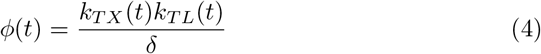

Thus the expression rate of the reporter *ϕ*(*t*) depends on both transcription and translation rates. Both of these rates are time varying and regulated in response to changing environmental conditions [Klumpp et al., 2009].

To model the growth and gene expression measurements from a population of cells we use the following equations,

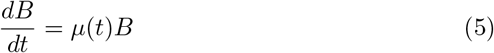

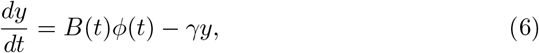

where *B* is a measure of sample biomass, *μ*(*t*) is the instantaneous relative growth rate, *y* is the intensity of reporter, *ϕ*(*t*) is the instantaneous expression rate, and *γ* is the reporter degradation rate. Here we have assumed that reporter intensity *y* = *Bp*, with *p* the protein concentration per biomass.

From these equations we wish to accurately and robustly estimate the growth rate *μ*(*t*) and the expression rate *ϕ*(*t*), given an estimate of the reporter degradation rate *γ*. Typically reporter proteins are stable and so in the following we will use *γ* = 0 [Andersen et al., 1998]. Reconstructing the functions *μ*(*t*) and *ϕ*(*t*) represents an inverse problem, which is underdetermined and ill-posed [Beck and Woodbury, 1998].

### Approximating dynamics with a Gaussian basis

In order to reduce the dimensionality of the problem we exploit prior knowledge of the functions *μ*(*t*) and *ϕ*(*t*) to construct a simple basis as follows. Expression rates and growth rates may be reasonably assumed to be strictly positive, and smooth on typical time scales of transcription and translation. We propose the following approximation, given a function *f*(*t*) that meets our assumptions,

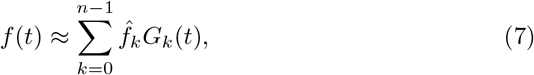

with,

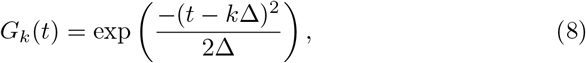

which represents a sum of *n* Gaussian curves *G_k_* of weight 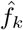 regularly spaced over time t at intervals Δ and with variance Δ. Here Δ determines the time scale of variation or smoothness of the representation of the function *f*(*t*). Choosing Δ greater than the sampling interval of the data makes the system over-determined and regularized in the sense that it is constrained to be smooth. In the following we choose Δ = 1 hour, the typical time scale of protein synthesis, which is larger than the usual sampling interval of 10-15 minutes.

### Dynamics estimation as an inverse problem

The model given in equations 5 and 6 represents the forward models of the inverse problems for reconstruction of *μ*(*t*) and *ϕ*(*t*). In practice the measurements used to estimate *B* and *y* are discrete, will contain background signal, and are subject to noise. The background signals *B*′ and *y*′ are typically estimated by measuring appropriate control samples containing no cells (*B*′) and cells with no reporter expression (*y*′). After subtracting these background measurements we are left with the noisy estimates 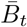 and 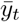. We then wish to parameterize the forward models given by equations 5 and 6 such that 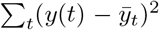 and 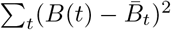 are minimized.

### Accurate estimation of growth rate dynamics

The inverse problem for reconstruction of the growth rate *μ*(*t*) can be stated as,

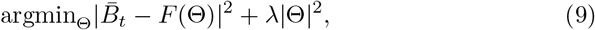

with,

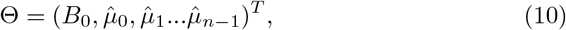

using *n* Gaussian basis functions to represent *μ*(*t*), and *F*(Θ) is the forward model. Since this problem is ill-posed we regularize using a Tikhonov penalty term. The hyperparameter λ may be chosen by various methods, here we use the L-curve [Hansen and O’Leary, 1993] (see Methods). This problem is a nonlinear least squares optimization, which we solve using the trust region reflective algorithm [Branch et al., 1999].

Figure 2 shows the result of fitting to simulated data generated from equations 5 and 6 using 100 randomly parameterized Gompertz growth models [Zwietering et al., 1990], and three different levels of measurement noise. We compare to the direct linear inversion method and show that our approach reduces mean squared error by more than 25-fold (*p* < 10^-32^, Welch’s T-test) (figure 2B). Further, our method is more robust to noise in early biomass measurements so that it correctly reconstructs the lag phase, which is missing from the linear inversion solution. Our method also reconstructs better the growth rate peak that can be used to separate exponential and stationary growth phases (figure 2C).

**Figure 1:**
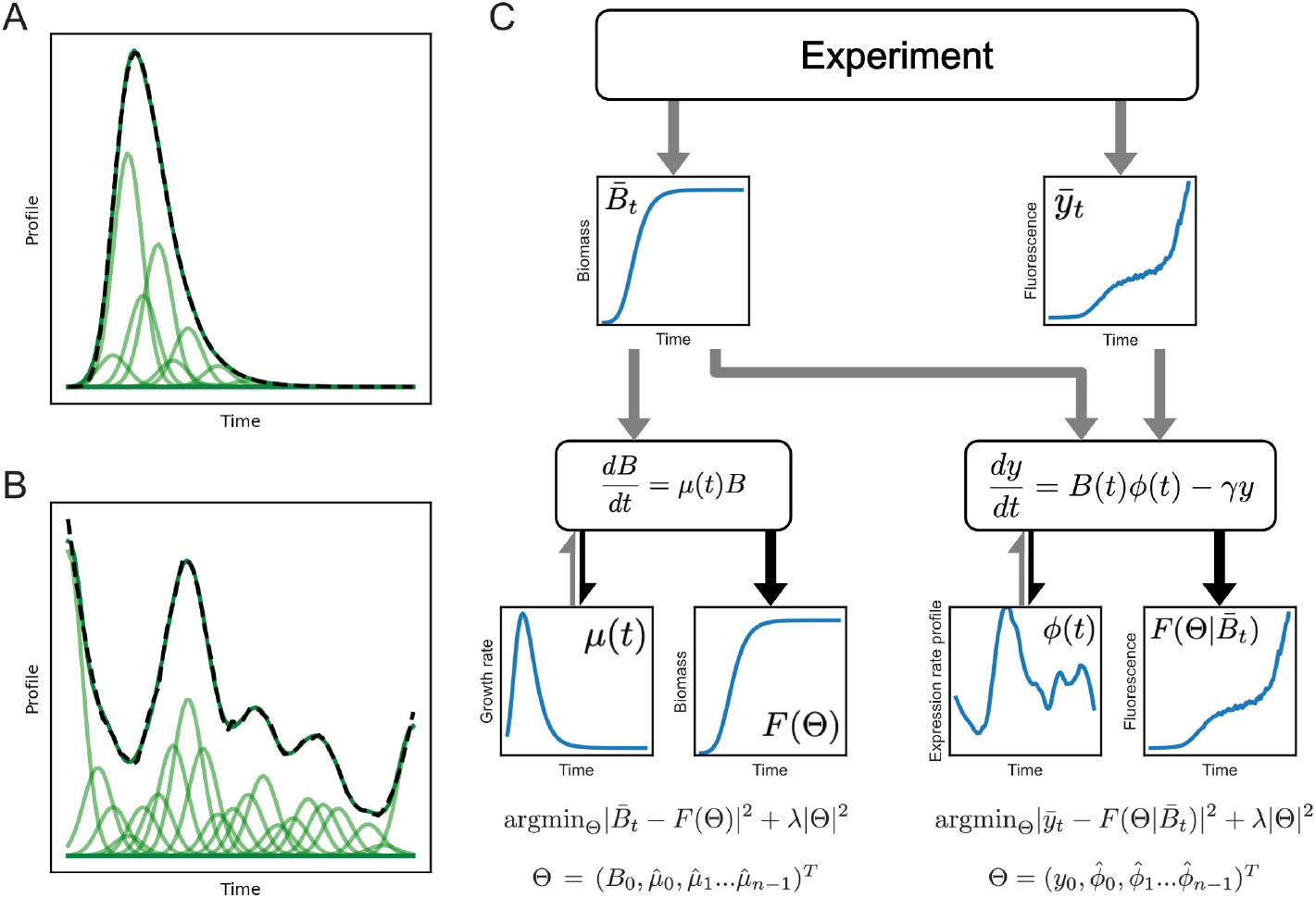
Overview of algorithm. Growth rate profiles (A, dashed black line) and gene expression rate profiles (B, dashed black line) can be approximated as a sum of Gaussian functions (A and B, dark green line). Individual basis functions are shown in light green. (C) Experimental time series data is used as inputs to differential equation forward models (*F*(Θ)) of growth (left) and gene expression (right). Each forward model also takes as input estimates of the corresponding dynamic rate profile Θ (formed from a Gaussian basis) and outputs simulated measurement time series. The final reconstruction is found by least squares minimization of the difference between the forward model and the data.

**Figure 2:**
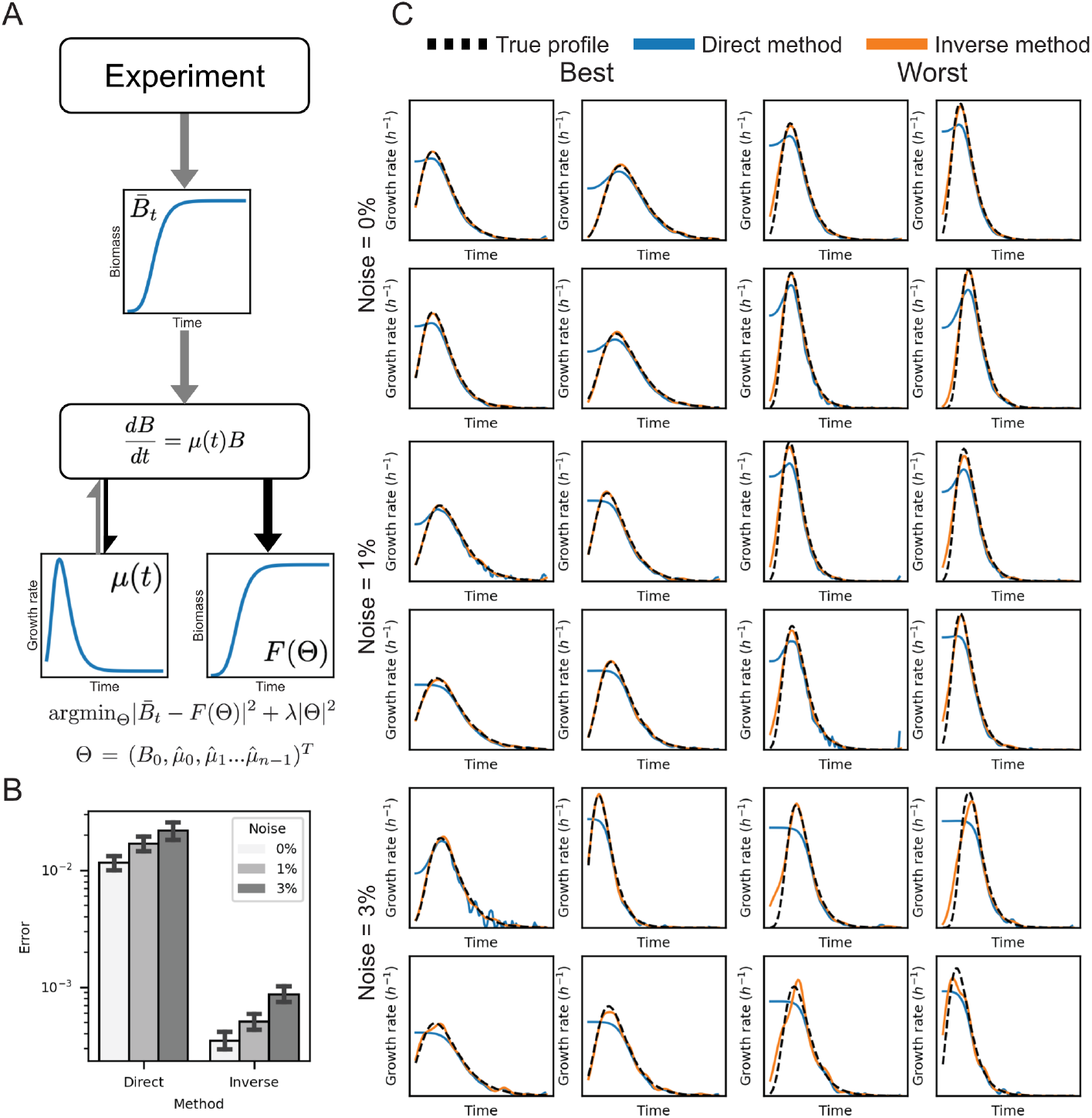
Reconstructing growth rate from noisy biomass measurements. (A) Overview of algorithm. Given a candidate growth rate profile *μ*(*t*), the forward model consists of a simple differential equation for biomass. This equation is solved numerically and the squared error of the resulting profile from the ex-perimental data is minimized. (B) Compared with the direct linear inversion method, reconstruction of simulated Gompertz growth rate profiles is significantly more accurate. (C) The best and worst resulting profiles at different noise levels. Direct linear inversion fails to accurately reconstruct lag phase growth.

### Accurate estimation of gene expression rate dynamics

Following equation 9 we wish to find the optimal parameters,

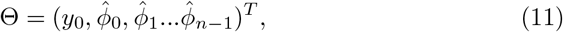

with the forward model conditional on the biomass 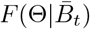. The problem is again a non-linear least squares optimization,

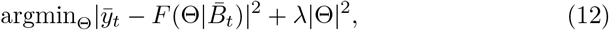

which we solve using the same numerical procedure as for growth rate.

To test this approach we generated 100 random gene expression rate profiles from smoothed log-normal random walks, with random Gompertz models for the biomass, and three measurement noise levels (see Methods). Again we compared to the direct linear inversion method and show that the mean squared error is more than 2-fold lower (*p* < 5 × 10^-6^, Welch’s T-test) (figure 3B). We also find that the direct method fails, even with no noise, on some profile shapes, producing extremely noisy solutions (figure 3C).

**Figure 3:**
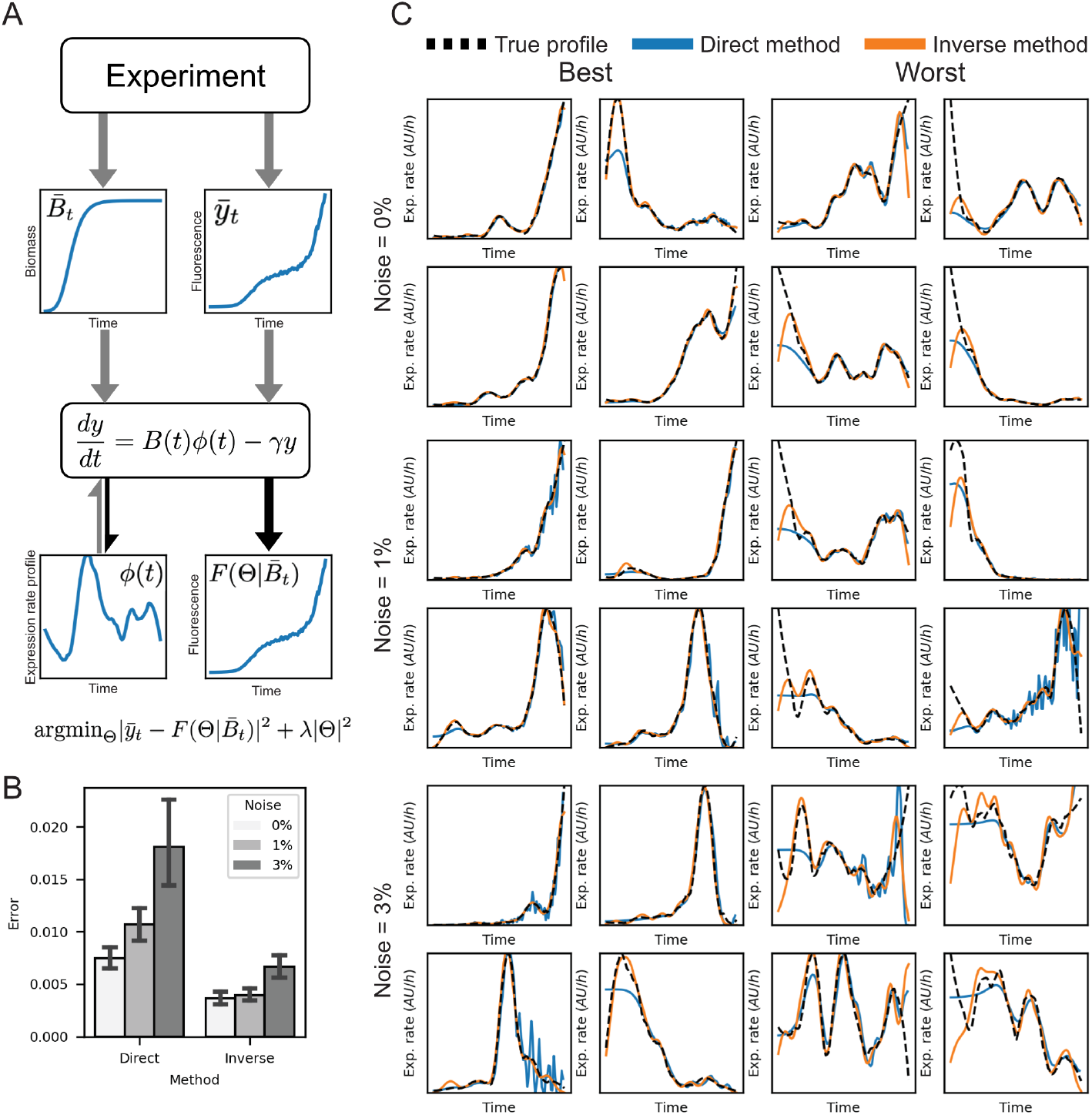
Reconstructing gene expression rate from noisy measurements. (A) Overview of algorithm. Given a candidate gene expression rate profile *ϕ*(*t*), and a background corrected biomass time series 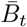 the forward model consists of a simple differential equation for reporter intensity. This equation is solved numerically and the squared error of the resulting profile from the background corrected reporter time series is minimized. (B) Compared with the direct linear inversion method, reconstruction of random gene expression rate profiles is significantly more accurate. (C) The four best and four worst resulting profiles at different noise levels. Direct linear inversion fails with very noisy solutions.

### Promoter-reporter fusions do not reflect intrinsic promoter transcription rate dynamics

Promoter-reporter fusions are often used to characterize the dynamic transcription rates of promoters in natural and synthetic genetic regulatory networks. However, the dynamics of reporter expression depend on transcription and translation, which both vary according to growth conditions. We reconstructed the growth and gene expression dynamics for *Escherichia coli* carrying a synthetic triple promoter-reporter fusion plasmid (figure 4A). This plasmid encoded for three different fluorescent reporters from genes driven by the same synthetic constitutive *σ*_70_ promoter [iGEM, 2006]. Measurement assays were performed in two different strains, with each growing on two different carbon sources [Yáñez Feliú et al., 2020]. Assays were performed using a 96-well microplate reader to measure fluorescence in the three channels as well as optical density as a proxy for biomass, with a sampling interval of 15 minutes.

**Figure 4:**
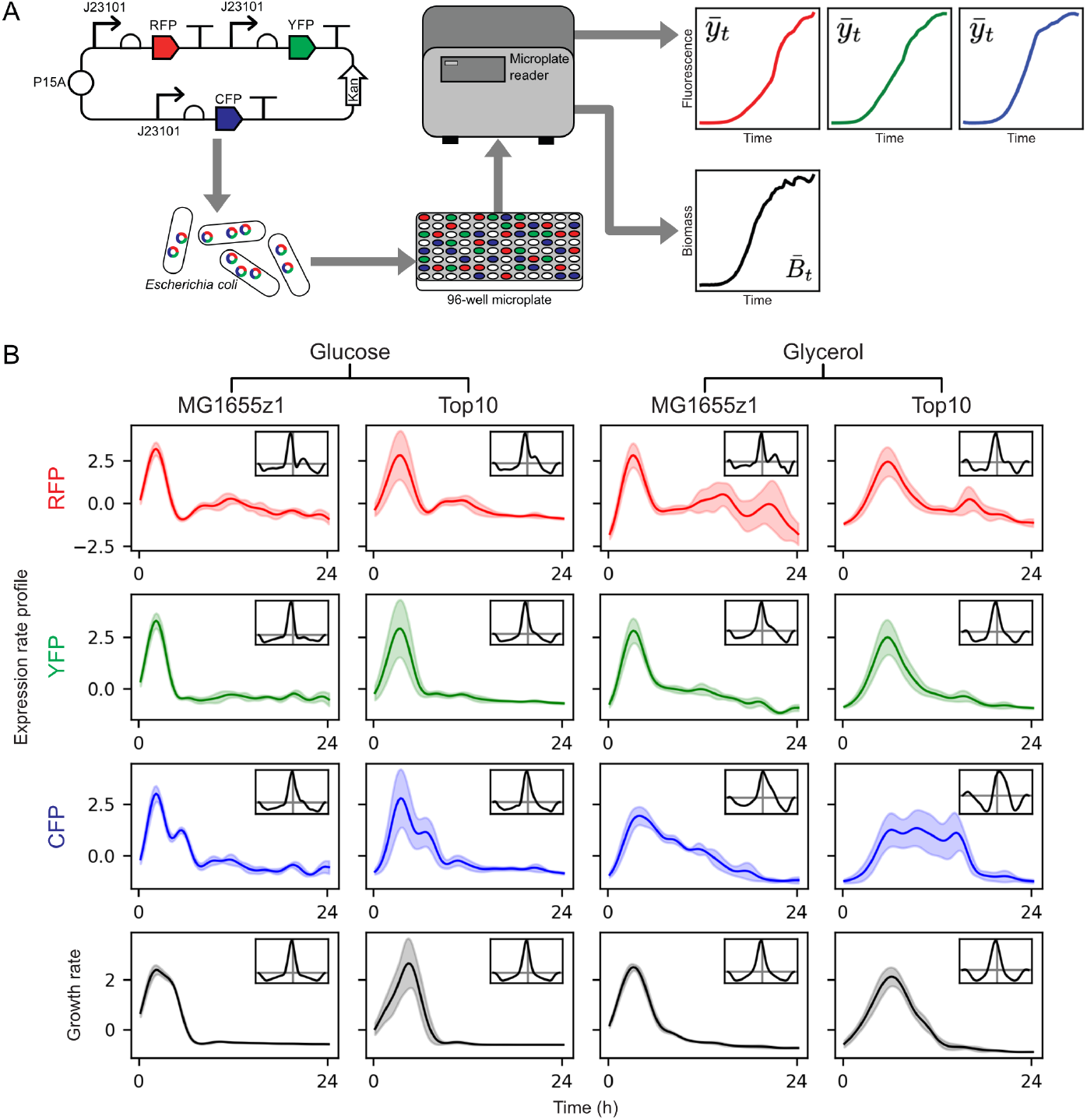
Reconstruction of gene expression and growth rate profiles from experimental data. (A) Experimental procedure. A plasmid encoding three promoterreporter fusion genes with the same promoter (J23101) was transformed into *E. coli*., which were inocculated into a 96-well microplate and measured in a microplate reader. The three reporter intensities and optical density as a measure of biomass were recorded approximately every 15 minutes. (B) The resulting gene expression rate profiles (top three rows) and growth rate profile (bottom row). Insets show cross-correlation with growth rate profile. Both growth conditions (carbon source glucose or glycerol, and strain MG1655z1 or Top10) and downstream reporter (RFP, YFP or CFP) change the dynamics of gene expression rate. Plots show mean and standard deviation of 30 replicates from 3 separate experiments.

Figure 4B shows the gene expression rate profiles for each fluorescent reporter gene (top three rows), and the growth rate for the corresponding samples (bottom row). Consistent with observed correlations between growth rate and gene expression rates [Klumpp et al., 2009], gene expression rates broadly peak at maximal growth rate. Effects on the qualitative form can also be seen from the cross-correlation of gene expression rates with growth rates (figure 4B, insets). Different carbon sources and host strains both impact on the dynamics of gene expression and growth rate as expected. However, we also see that the common promoter sequence contained in each fusion gene does not generate the same gene expression profile. This suggests that either the context of the promoter - its position in the plasmid, orientation, flanking sequences - affect the dynamics of transcription, or that other factors such as translation efficiency also vary over time and are different for the different genes. In either case, it is clear that promoter-reporter fusion gene expression rate profiles should not in general be interpreted as indicative of intrinsic promoter transcription rate dynamics.

## Discussion

We have demonstrated an inverse problems approach to reconstructing dynamic gene expression rate and growth rate dynamic profiles from noisy kinetic measurement data. We compared our method to the current state-of-the-art algorithm, direct linear inversion [Zulkower et al., 2015]. Our approach reduced the mean squared error of reconstructions from simulated data of growth rate by 25-fold and gene expression rate by more than two-fold. Further, we were able to reconstruct features of both growth rate and gene expression rate profiles that were not apparent from the direct linear inversion method.

Accurate reconstruction of dynamic gene expression and growth rate profiles is essential for characterization of synthetic genetic circuits and for inference of gene regulatory interactions in natural networks [Berthoumieux et al., 2013, Shin et al., 2020]. Improving genetic circuit characterization we will feed modelling and design tools with better data. The reconstruction of lag and exponential phase patterns in growth profiles is key for growth phase segmentation.

Typically measurements of promoter-reporter fusions are used as a proxy for transcription rates under various levels of transcription factors, external signals, and other determinants of gene expression [Nielsen et al., 2016, Tas et al., 2021, Rudge et al., 2016, Shin et al., 2020]. However, applying our method to multiple reporter fusions with the same promoter, we show that gene expression rate profiles should not be taken as indicative of intrinsic characteristics of promoters, since they are affected by both the promoter and the downstream reporter, as well as potentially other factors. This also indicates that ratiometric techniques where a standard transcriptional unit is taken as expression reference [Nielsen et al., 2016, Rudge et al., 2016, Kelly et al., 2009] are not valid for dynamic gene expression rates.

## Acknowledgements

G.V. was supported by a scholarship from the Institute for Biological and Medical Engineering, Pontificia Universidad Católica de Chile. G.V., C.V. and T.J.R. was supported by ANID PIA Anillo ACT192015 and ANID Fondecyt Regular 1211598. G.V. and T.J.R. were supported by ANID Fondecyt Iniciación 11161046. The authors thank Carlos Sing Long and Simon Arridge for helpful comments and discussions.

## Author Contributions

Conceptualization, T.J.R.; Methodology, G.V., C.V., and T.J.R.; Investigation, G.V., C.V., and T.J.R.; Writing – Original Draft, G.V., C.V., and T.J.R.; Writing – Review Editing, G.V. C.V., and T.J.R.; Funding Acquisition, T.J.R.; Resources, T.J.R.; Supervision, T.J.R.

## Figure Legends

**Figure 1**. Overview of algorithm. Growth rate profiles (A, dashed black line) and gene expression rate profiles (B, dashed black line) can be approximated as a sum of Gaussian functions (A and B, dark green line). Individual basis functions are shown in light green. (C) Experimental time series data is used as inputs to differential equation forward models (*F*(Θ)) of growth (left) and gene expression (right). Each forward model also takes as input estimates of the corresponding dynamic rate profile Θ (formed from a Gaussian basis) and outputs simulated measurement time series. The final reconstruction is found by least squares minimization of the difference between the forward model and the data.

**Figure 2**. Reconstructing gene expression rate from noisy biomass measurements. (A) Overview of algorithm. Given a candidate gene expression rate profile *ϕ*(*t*), and a background corrected biomass time series 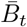 the forward model consists of a simple differential equation for reporter intensity. This equation is solved numerically and the squared error of the resulting profile from the background corrected reporter time series is minimized. (B) Compared with the direct linear inversion method, reconstruction of random gene expression rate profiles is significantly more accurate. (C) The four best and four worst resulting profiles at different noise levels. Direct linear inversion fails with very noisy solutions.

**Figure 3**. Reconstructing gene expression rate from noisy measurements. (A) Overview of algorithm. Given a candidate gene expression rate profile *ϕ*(*t*), and a background corrected biomass time series 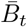 the forward model consists of a simple differential equation for reporter intensity. This equation is solved numerically and the squared error of the resulting profile from the background corrected reporter time series is minimized. (B) Compared with the direct linear inversion method, reconstruction of random gene expression rate profiles is significantly more accurate. (C) The four best and four worst resulting profiles at different noise levels. Direct linear inversion fails with very noisy solutions.

**Figure 4**. Reconstruction of gene expression and growth rate profiles from experimental data. (A) Experimental procedure. A plasmid encoding three promoter-reporter fusion genes with the same promoter (J23101) was transformed into *E. coli*., which were inocculated into a 96-well microplate and measured in a microplate reader. The three reporter intensities and optical density as a measure of biomass were recorded approximately every 15 minutes. (B) The resulting gene expression rate profiles (top three rows) and growth rate profile (bottom row). Insets show cross-correlation with growth rate profile. Both growth conditions (carbon source glucose or glycerol, and strain MG1655z1 or Top10) and downstream reporter (RFP, YFP or CFP) change the dynamics of gene expression rate. Plots show mean and standard deviation of 30 replicates from 3 separate experiments.

## Methods

### RESOURCE AVAILABILITY

#### Lead Contact

Further information and requests for resources and reagents should be directed to and will be fulfilled by the Lead Contact, Timothy J. Rudge.

#### Materials Availability

This study did not generate new materials. The triple promoter-reporter fusion plasmids from [Yáñez Feliú et al., 2020] were used to reconstruct growth and gene expression dynamics.

#### Data and Code Availability

The synthetic and experimental data generated during this study are available in Flapjack (http://flapjack.rudge-lab.org) All original code is open and accessible at https://github.com/synbiouc/flapjack_api.

### EXPERIMENTAL MODELS AND SUBJECT DETAILS

Two *E. coli* strains were used in this study: TOP10 and MG1655 Z1 malE.

### METHOD DETAILS

#### Experimental Methods

Here we describe the physical experiments reported in this study.

#### Kinetic gene expression and growth assays

To assay gene expression we cultured bacteria containing the triple reporter plasmid pAAA (figure 4A) described in [Yáñez Feliú et al., 2020], in two different strains and two different carbon sources.

#### Experiment

Monoclonal colonies of bacteria TOP10 or MG1655 malE transformed with plasmid pAAA were picked and cultured for 14-15 hours overnight in M9 media with 50 *μ*g/mL kanamycin, 0.2% w/v casaminoacids and 0.4% w/v glucosa or 0.4% v/v glicerol. Overnight cultures were diluted 1000 times in 2 mL tubes. All the tubes were filled with 1996 *μ*L of fresh M9 media, 2 *μ*L of kanamycin and 2 *μ*L of the bacteria liquid culture obtaining a final volume of 2 mL. In each well of a 96 well plate were added 200 *μ*L, 4 wells with M9 media with the proper carbon source and kanamycin, 4 wells with non transformed bacteria of the same strain and 10 wells of bacteria transformed with pAAA to analyze from the previously prepared 2 mL tubes. Optical density and fluorescence in three channels (RFP, YFP and CFP) were measured approximately every 15 minutes for 24 hours in a Synergy HTX plate reader with Gen5 software. Each assay was repeated on 3 different days, and each 96 well plate contained experiments with the same carbon source and strain following the methods in [Yáñez Feliú et al., 2020].

#### Computational Methods

All computation was performed using Flapjack [Yáñez Feliú et al., 2020] and Python [Van Rossum and Drake, 2009], Numpy [Harris et al., 2020], Scipy [Virtanen et al., 2020], Pandas [McKinney, 2010], Matplotlib [Hunter, 2007] and Plotly [Plotly Technologies Inc., 2015].

#### Web-based software implementation

We extended Flapjack [Yáñez Feliú et al., 2020] (http://flapjack.rudge-lab.org) to compute gene expression rate and growth rate profiles using the methods described above. Flapjack is a systems and synthetic biology data storage and analysis tool, built as a web app that provides a user-friendly web interface and a REST and web socket API. The system allows upload of kinetic gene expression data from a variety of sources, and links it to metadata about experimental conditions. These data may then be queried and filtered, and used to reconstruct gene expression and growth rates. Flapjack automatically subtracts background signal from both reporter and biomass measurements, taking the average of control samples (untransformed cells or media with no cells) at each time point.

#### Random profile generation

In order to generate a range of gene expression rate profiles, with minimal assumptions about their form, we generate log-normal random walks as,

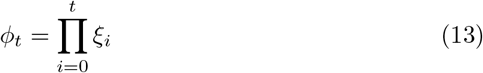

with log(*ξ_i_*) ~ *N*(0, *σ*^2^), and *σ*^2^ = 0.25. The profiles *ϕ_t_* were then smoothed using a second-order Savitzky-Golay filter [Savitzky and Golay, 1964] with window size 21, and normalized to [0, 1].

To generate random growth rate profiles we used the Gompertz equation,

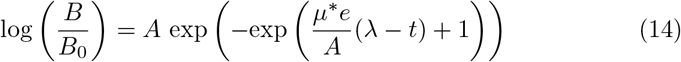

with *μ*\* the maximal growth rate uniformly distributed on [0.5, 1] per hour, λ the lag phase length uniformly distributed in [0, 4] hours, *A* = log(*B*\*/*B*_0_), where the maximal biomass *B*\* = 1 and minimum biomass *B*_0_ = 0.01. The growth rate profile implied by this equation is given by,

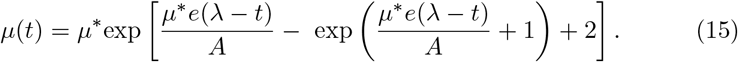

#### Simulation of kinetic gene expression and biomass data

Equations 5 and 6 were solved using the forward Euler integration scheme with time step Δ*t* = 2.4 × 10^-2^ hours for a period of 24 hours. Noise and background were added according to the following equations with *B*′ = 0.1 and *y*′ = 0.1,

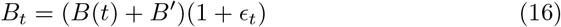

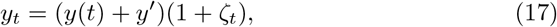

where *ϵ_t_* and *ζ_t_* are uncorrelated white noise with variance *σ*^2^, due to the measurement process. Simulated measurements were then uploaded to Flapjack [Yáñez Feliú et al., 2020], and analyzed using the API via Python.

#### Choice of regularization parameter

The Tikhonov regularization parameter λ was chosen using the L-curve. The norm of the solution (Θ) was plotted against the least-squares error for a range of λ and the value at corner of the curve was chosen.

## References

[Aïchaoui et al., 2012] Aïchaoui, L., Jules, M., Le Chat, L., Aymerich, S., Fromion, V., and Goelzer, A. (2012). Basylica: a tool for automatic processing of a bacterial live cell array. Bioinformatics, 28(20):2705–2706.

[Andersen et al., 1998] Andersen, J. B., Sternberg, C., Poulsen, L. K., Bjørn, S. P., Givskov, M., and Molin, S. (1998). New unstable variants of green fluorescent protein for studies of transient gene expression in bacteria. Applied and environmental microbiology, 64(6):2240–2246.

[Beal et al., 2020] Beal, J., Farny, N. G., Haddock-Angelli, T., Selvarajah, V., Baldwin, G. S., Buckley-Taylor, R., Gershater, M., Kiga, D., Marken, J., Sanchania, V., et al. (2020). Robust estimation of bacterial cell count from optical density. Communications biology, 3(1):1–29.

[Beal et al., 2018] Beal, J., Haddock-Angelli, T., Baldwin, G., Gershater, M., Dwijayanti, A., Storch, M., De Mora, K., Lizarazo, M., Rettberg, R., and with the iGEM Interlab Study Contributors (2018). Quantification of bacterial fluorescence using independent calibrants. PloS one, 13(6):e0199432.

[Beck and Woodbury, 1998] Beck, J. V. and Woodbury, K. A. (1998). Inverse problems and parameter estimation: integration of measurements and analysis. Measurement Science and Technology, 9(6):839.

[Berthoumieux et al., 2013] Berthoumieux, S., De Jong, H., Baptist, G., Pinel, C., Ranquet, C., Ropers, D., and Geiselmann, J. (2013). Shared control of gene expression in bacteria by transcription factors and global physiology of the cell. Molecular systems biology, 9(1):634.

[Branch et al., 1999] Branch, M. A., Coleman, T. F., and Li, Y. (1999). A subspace, interior, and conjugate gradient method for large-scale bound-constrained minimization problems. SIAM Journal on Scientific Computing, 21(1):1–23.

[De Jong et al., 2010] De Jong, H., Ranquet, C., Ropers, D., Pinel, C., and Geiselmann, J. (2010). Experimental and computational validation of models of fluorescent and luminescent reporter genes in bacteria. BMC systems biology, 4(1):1–17.

[Hansen and O’Leary, 1993] Hansen, P. C. and O’Leary, D. P. (1993). The use of the l-curve in the regularization of discrete ill-posed problems. SIAM journal on scientific computing, 14(6):1487–1503.

[Harris et al., 2020] Harris, C. R., Millman, K. J., van der Walt, S. J., Gommers, R., Virtanen, P., Cournapeau, D., Wieser, E., Taylor, J., Berg, S., Smith, N. J., Kern, R., Picus, M., Hoyer, S., van Kerkwijk, M. H., Brett, M., Haldane, A., Río, J. F. D., Wiebe, M., Peterson, P., Gérard-Marchant, P., Sheppard, K., Reddy, T., Weckesser, W., Abbasi, H., Gohlke, C., and Oliphant, T. E. (2020). Array programming with numpy. Nature, 585(7825):357–362.

[Hunter, 2007] Hunter, J. D. (2007). Matplotlib: A 2d graphics environment. Computing in Science & Engineering, 9(3):90–95.

[iGEM, 2006] iGEM (2006). Registry of standard biological parts. http://parts.igem.org/Promoters/Catalog/Anderson (Accessed on 2021-03-15).

[Kalir et al., 2001] Kalir, S., McClure, J., Pabbaraju, K., Southward, C., Ronen, M., Leibler, S., Surette, M., and Alon, U. (2001). Ordering genes in a flagella pathway by analysis of expression kinetics from living bacteria. Science, 292(5524):2080–2083.

[Kelly et al., 2009] Kelly, J. R., Rubin, A. J., Davis, J. H., Ajo-Franklin, C. M., Cumbers, J., Czar, M. J., de Mora, K., Glieberman, A. L., Monie, D. D., and Endy, D. (2009). Measuring the activity of biobrick promoters using an in vivo reference standard. Journal of Biological Engineering, 3(1):4–4.

[Klumpp et al., 2009] Klumpp, S., Zhang, Z., and Hwa, T. (2009). Growth rate-dependent global effects on gene expression in bacteria. Cell, 139(7):1366–1375.

[Lichten et al., 2014] Lichten, C. A., White, R., Clark, I. B., and Swain, P. S. (2014). Unmixing of fluorescence spectra to resolve quantitative time-series measurements of gene expression in plate readers. BMC biotechnology, 14(1):1–11.

[McKinney, 2010] McKinney, W. (2010). Data structures for statistical computing in python. In Proceedings of the 9th Python in Science Conference,pages 56–61.

[Nielsen et al., 2016] Nielsen, A. A., Der, B. S., Shin, J., Vaidyanathan, P., Paralanov, V., Strychalski, E. A., Ross, D., Densmore, D., and Voigt, C. A. (2016). Genetic circuit design automation. Science, 352(6281).

[Nuñez et al., 2017] Nuñez, I., Matute, T., Herrera, R., Keymer, J., Marzullo, T., Rudge, T., and Federici, F. (2017). Low cost and open source multifluorescence imaging system for teaching and research in biology and bioengineering. PLOS ONE, 12(11):1–21.

[Plotly Technologies Inc., 2015] Plotly Technologies Inc. (2015). Collaborative data science. https://plot.ly (Accessed on 2021-01-11).

[Ronen et al., 2002] Ronen, M., Rosenberg, R., Shraiman, B. I., and Alon, U. (2002). Assigning numbers to the arrows: parameterizing a gene regulation network by using accurate expression kinetics. Proceedings of the national academy of sciences, 99(16):10555–10560.

[Rudge et al., 2016] Rudge, T. J., Brown, J. R., Federici, F., Dalchau, N., Phillips, A., Ajioka, J. W., and Haseloff, J. (2016). Characterization of intrinsic properties of promoters. ACS synthetic biology, 5(1):89–98.

[Savitzky and Golay, 1964] Savitzky, A. and Golay, M. J. (1964). Smoothing and differentiation of data by simplified least squares procedures. Analytical chemistry, 36(8):1627–1639.

[Shin et al., 2020] Shin, J., Zhang, S., Der, B. S., Nielsen, A. A., and Voigt, C. A. (2020). Programming escherichia coli to function as a digital display. Molecular systems biology, 16(3):e9401.

[Tas et al., 2021] Tas, H., Grozinger, L., Stoof, R., de Lorenzo, V., and Goñi-Moreno, Á. (2021). Contextual dependencies expand the re-usability of genetic inverters. Nature communications, 12(1):1–9.

[Van Rossum and Drake, 2009] Van Rossum, G. and Drake, F. L. (2009). Python 3 Reference Manual. CreateSpace, Scotts Valley, CA.

[Virtanen et al., 2020] Virtanen, P., Gommers, R., Oliphant, T. E., Haberland, M., Reddy, T., Cournapeau, D., Burovski, E., Peterson, P., Weckesser, W., Bright, J., van der Walt, S. J., Brett, M., Wilson, J., Millman, K. J., Mayorov, N., Nelson, A. R. J., Jones, E., Kern, R., Larson, E., Carey, C. J., Polat, I., Feng, Y., Moore, E. W., VanderPlas, J., Laxalde, D., Perktold, J., Cimrman, R., Henriksen, I., Quintero, E. A., Harris, C. R., Archibald, A. M., Ribeiro, A. H., Pedregosa, F., van Mulbregt, P., and SciPy 1.0 Contributors (2020). SciPy 1.0: Fundamental Algorithms for Scientific Computing in Python. Nature Methods, 17:261–272.

[Yáñez Feliú et al., 2020] Yáñez Feliú, G., Earle Gómez, B., Codoceo Berrocal, V., Muñoz Silva, M., Nuñez, I. N., Matute, T. F., Arce Medina, A., Vidal, G., Vidal Céspedes, C., Dahlin, J., Federici, F., and Rudge, T. J. (2020). Flapjack: Data management and analysis for genetic circuit characterization. ACS Synthetic Biology, 0(0):null.

[Zulkower et al., 2015] Zulkower, V., Page, M., Ropers, D., Geiselmann, J., and de Jong, H. (2015). Robust reconstruction of gene expression profiles from reporter gene data using linear inversion. Bioinformatics, 31(12):71–79.

[Zwietering et al., 1990] Zwietering, M. H., Jongenburger, I., Rombouts, F. M., and van’t Riet, K. (1990). Modeling of the bacterial growth curve. Applied and Environmental Microbiology, 56(6):1875–1881.

